# A massively parallel *in vivo* assay of TdT mutants yields variants with altered nucleotide insertion biases

**DOI:** 10.1101/2024.06.11.598561

**Authors:** Courtney K. Carlson, Theresa B. Loveless, Marija Milisavljevic, Patrick I. Kelly, Jeremy H. Mills, Keith E. J. Tyo, Chang C. Liu

## Abstract

Terminal deoxynucleotidyl transferase (TdT) is a unique DNA polymerase capable of template-independent extension of DNA with random nucleotides. TdT’s *de novo* DNA synthesis ability has found utility in DNA recording, DNA data storage, oligonucleotide synthesis, and nucleic acid labeling, but TdT’s intrinsic nucleotide biases limit its versatility in such applications. Here, we describe a multiplexed assay for profiling and engineering the bias and overall activity of TdT variants in high throughput. In our assay, a library of TdTs is encoded next to a CRISPR-Cas9 target site in HEK293T cells. Upon transfection of Cas9 and sgRNA, the target site is cut, allowing TdT to intercept the double strand break and add nucleotides. Each resulting insertion is sequenced alongside the identity of the TdT variant that generated it. Using this assay, 25,623 unique TdT variants, constructed by site-saturation mutagenesis at strategic positions, were profiled. This resulted in the isolation of several altered-bias TdTs that expanded the capabilities of our TdT-based DNA recording system, Cell History Recording by Ordered Insertion (CHYRON), by increasing the information density of recording through an unbiased TdT and achieving dual-channel recording of two distinct inducers (hypoxia and Wnt) through two differently biased TdTs. Select TdT variants were also tested *in vitro*, revealing concordance between each variant’s *in vitro* bias and the *in vivo* bias determined from the multiplexed high throughput assay. Overall, our work, and the multiplex assay it features, should support the continued development of TdT-based DNA recorders, *in vitro* applications of TdT, and further study of the biology of TdT.

## Introduction

Terminal deoxynucleotidyl transferase (TdT) is a DNA polymerase that extends DNA in a template-independent manner through the polymerization of random nucleotides^1,2^. The unique ability of TdT to carry out *de novo* DNA synthesis has been leveraged in numerous applications. For some applications such as the progressive generation of high-information genetic barcodes *in vivo* for deep lineage tracing^3,4^ or the synthesis of aptamer libraries^5^, high randomness in the composition of TdT-generated sequences is desirable. For other applications such as DNA data storage^6–9^, enzymatic DNA synthesis^10–13^, nucleic acid end-labeling^14–18^, and certain examples in biosensing^19–26^, randomness is a liability to be reined in. An obvious way to shape the composition of TdT-synthesized sequences is by pre-defining the pool of nucleotides provided^14,15,19–27^. However, such control is only available *in vitro* and is far from total, given the strength of TdT’s intrinsic biases^2,28^. A broad range of TdT-based applications could benefit from engineered TdTs whose intrinsic biases and activities are modified.

In the present study, we developed a sequencing-based multiplexed assay^29–35^ that enabled pooled screening of tens of thousands of TdT variants in HEK293T cells. Each TdT’s activity and nucleotide biases were measured by “next-generation” sequencing (NGS) of the identity of the TdT variant alongside outcomes of TdT-mediated insertional mutagenesis at a CRISPR target site. As input to the assay, we made 3 site saturation mutagenesis libraries of TdT containing 27,783 possible amino acid sequences. 92.2% of these TdT variants were sufficiently sampled to determine nucleotide bias (≥447 reads for each variant, see Methods). To demonstrate the reliability of the pooled assay results, we selected 74 TdT variants to assay individually and found that the insertion mutation outcomes profiled in the pooled assay matched those determined in the individual assays. In addition, we explored whether TdT nucleotide biases determined *in vivo* were reproducible *in vitro* and observed general agreement. Finally, we demonstrated the utility of novel TdTs identified in our assay by using three variants with altered biases in our established DNA recorder, CHYRON^36^. A TdT variant whose nucleotide preferences were balanced between A/T and G/C insertions allowed for higher information density recording in CHYRON, while two other TdT variants, one with strong A/T bias (86% A/T) and the other with strong G/C bias (28% A/T), allowed for dual-channel recording of two distinct biological inputs. Taken together, the multiplexed nature of our TdT screen, the reproducibility of its results both *in vivo* and *in vitro*, and the immediate utility of altered-bias TdT variants in DNA recording should enable new applications for engineered TdTs.

## Results

### A library of TdT mutants yielded active TdT variants with novel biases

Wild-type TdTs have an intrinsic bias for incorporating Gs and Cs^2^, which has been observed both *in vitro*^37^ and *in vivo*^38–41^. However, when TdT-based DNA recording strategies such as CHYRON^36^ and DARLIN^3^ are used to generate random barcodes inside cells for lineage tracing, an unbiased TdT is desirable to maximize the Shannon entropy of each added nucleotide and minimize the chance that independent cell lineages will generate the same barcode. On the other hand, when TdT-based DNA recorders are used to document the history of biological signals, TdTs with strong biases may instead be desirable, as the pattern of inserted nucleotides could be used to infer which TdT created the insertions, enabling recording of biological signals that can be tied to the expression of different TdTs. These goals in DNA recording, and other goals for TdT-based DNA technologies, motivated us to engineer novel TdTs and to develop a general strategy to do so.

To manipulate the bias of TdT, three NNK-based site-saturation mutagenesis libraries were constructed, each targeting three residues that we hypothesized would affect nucleotide preferences (Figures 1a,b). Two of the libraries targeted residues in a 16 amino acid lariat-like loop that is unique to TdT and known to be responsible for its template-independent activity (Loop1)^2^. We suspected Loop1 could influence dNTP selection because it is found in various conformations in the available crystal structures of TdT, suggesting it may play a dynamic role in polymerization^42^. In one library (Library 1), we randomized three residues at the base of Loop1 (D399, H400, F401) because they could feasibly alter the range of motion of the loop. In another library (Library 2), residues in Loop1 that we hypothesized could move into contact with the incoming dNTP (D396, L398, D399) were chosen for randomization. Aside from Loop1, we also mutagenized residues R454, E457, R461, which are located within 6 Å of the bound nucleotide during polymerization and point toward the nucleobase^2^, potentially influencing nucleotide choice (Library 3). In total, the three libraries represented 27,783 possible protein sequences (including stop codons) encoded by 98,304 mutant TdT DNA sequences, from which we hoped to find active TdT variants with altered biases.

**Figure 1.**
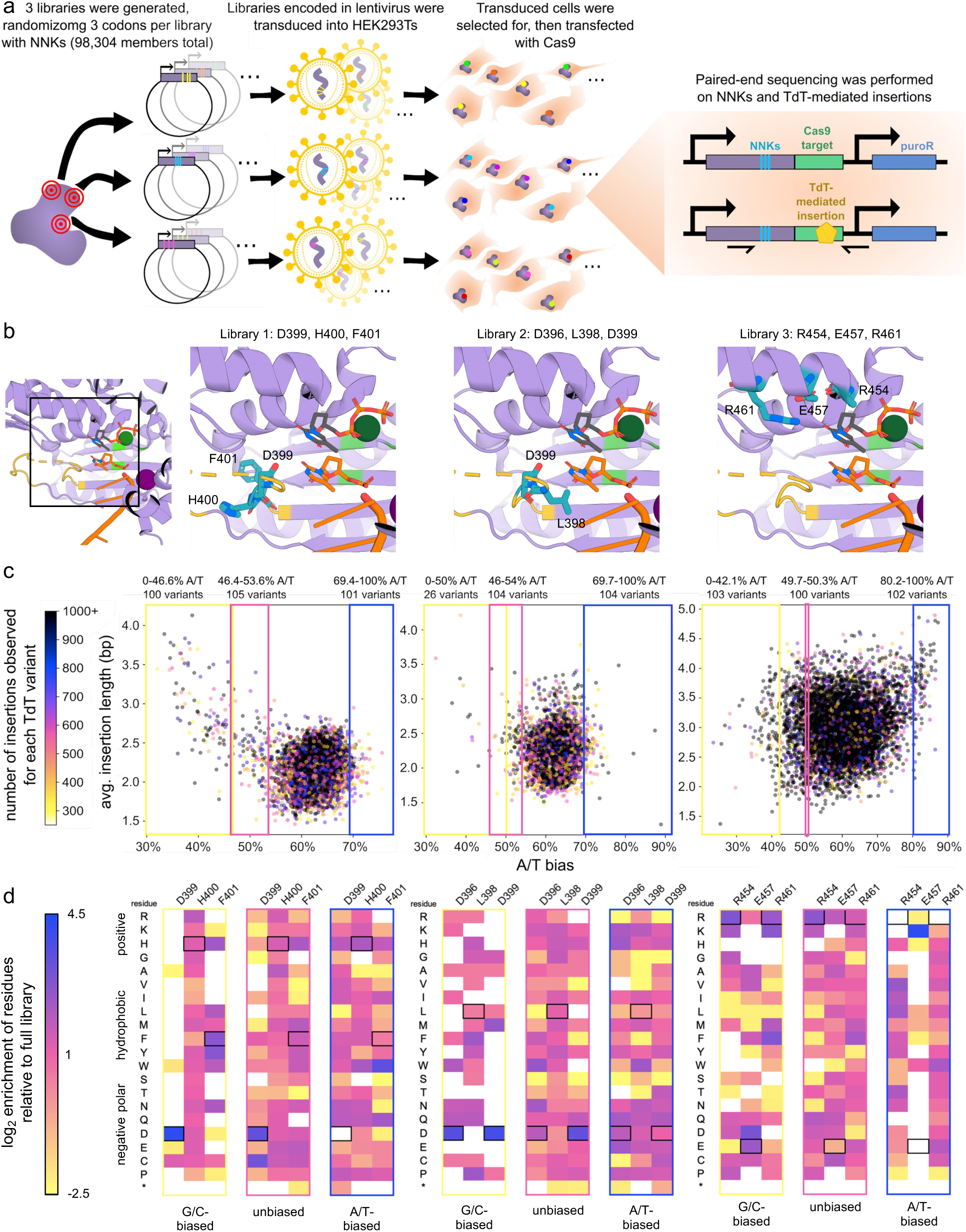
A massively parallel assay was conducted on 3 site saturation mutagenesis libraries of TdT. **a,** Workflow for creating the saturation mutagenesis libraries encoded in lentivirus and assaying the library members in HEK293Ts. Each library member was expressed in hundreds to thousands of cells, yielding many independent instances of each TdT variant. **b,** Crystal structures of TdT, highlighting the residues that were targeted for saturation mutagenesis in each library. Zoomed-out view (left-most) shows TdT bound to a single-stranded DNA primer (single helix colored orange, with 3’ terminal nucleotide shown in greater detail with red and blue indicating oxygen and nitrogen, respectively) and an incoming ddTTP (thymidine with a gray, red, and blue base moiety and orange and red triphosphate group). Loop1, which is disordered, is colored in yellow. Positions of the catalytic aspartates are indicated in lime green. Dark green and plum spheres are divalent magnesium and sodium, respectively. Zoomed-in views of the active site highlight the mutagenized residues in teal for each library, with red and blue indicating oxygen and nitrogen, in licorice representation. **c,** Scatterplots depicting the combinations of A/T biases and average insertion lengths among all active TdT variants (insertion fraction ≥ 0.7) that yielded at least 250 insertions in the high-throughput screen. Single points represent individual TdT variants. Color-coding for each point reflects the number of insertion sequences analyzed for each variant. Yellow, pink, and blue boxes demarcate bins of the ∼100 most G/C-biased, unbiased, and A/T-biased TdTs in each library. **d,** Composition of residues at the randomized positions within the G/C-biased, unbiased, and A/T-biased bins indicated in c. Heatmaps show the enrichment of each amino acid moiety, relative to the full library of variants that were sequenced deeply enough to have a chance of passing the activity and insertion cutoff filters (Methods, Supplementary Figure 1). White cells indicate residues that were absent in the bins (infinitely de-enriched). Black boxes outline the wildtype amino acids.

To assay the function of TdT libraries of this large size, we developed and performed a multiplexed screen (Figure 1a). Because the phenotype we aimed to achieve (*i.e.*, altered nucleotide insertion preferences) could be read out *via* NGS and because the target site for TdT-mediated insertions could be user-directed by a Cas9-generated double strand break (DSB) in mammalian cells^4,36,43^, we encoded TdT variants next to a CRISPR target locus at which TdT-mediated insertion would occur. This enabled us to read out the identity of a given TdT variant alongside the product of its action *via* NGS. Lentivirus populations encoding our three TdT libraries next to a CRISPR compatible target locus and a puromycin resistance cassette were made and used to transduce HEK293T cells at a multiplicity of infection (MOI) of 0.3 such that most transduced cells expressed only one mutant TdT (86% of cells that are infected receive only one virus particle)^44,45^. Successfully transduced cells were selected using puromycin. Library construction steps and the total numbers of lentiviral particles and cells used were scaled such that almost every unique TdT variant would be represented in multiple cells, allowing many independent trials of each variant’s insertion activity (see Methods).

The HEK293T cell population of TdT variants after lentivirus transduction became the starting point for assaying TdT activities in multiplex. To initiate TdT action, Cas9 and an sgRNA for the target locus immediately downstream of the TdT gene were transiently transfected into cells (Figure 1a). Cells were allowed to grow for 3 days during which each cell’s TdT variant had an opportunity to generate insertions at the Cas9-induced DSB. At the end of the experiment, we used paired-end Illumina sequencing to determine the exact insertions that each cell’s TdT variant made. This was done by using primers that amplified the diversified region of each TdT variant at one of the paired ends and the insertion sequence generated at the other paired end, effectively yielding a list of insertion sequences and the identity of the TdT variant that produced each. These data were analyzed to find TdT variants that have both high activity and altered biases.

When TdT is inactive, we predominantly observed deletions at the Cas9 target locus such that the “insertion fraction,” defined as (reads containing an insertion) / (reads containing any indel), was 0.27. We determined this value by co-expressing Cas9 with a catalytically disabled TdT (Supplementary Figure 2a). In contrast, for wild-type TdT, we observed an insertion fraction of 0.83. We therefore surmised that an insertion fraction exceeding ∼0.27 signifies that a given

TdT variant has at least some activity. A histogram of all TdT variants’ insertion fractions from our multiplexed assay exhibited a bimodal distribution (Supplementary Figure 2b-d), with a high insertion fraction cluster in the range of 0.7-1. Since we were interested in highly active TdTs for downstream applications, we chose a 0.7 insertion fraction as the cutoff for TdT variants we considered active. For bias, we evaluated the specific sequences of the insertions generated by active TdT variants. To confidently detect altered bias, evaluation of a few insertion sequences would be insufficient, because each TdT-generated insertion is short, averaging only a few nucleotides. Therefore, we only evaluated the biases of active TdT variants for which >250 insertion sequence reads were observed (see Methods). With both filters for activity (insertion fraction > 0.7) and insertion sequence count applied, 7193, 6640, and 6330 unique TdT variants from libraries 1, 2, and 3, respectively, remained, providing bias data on 77.7%, 71.7%, and 68.4% of all possible TdTs in libraries 1, 2, and 3, respectively. Plotting the average composition (A/T content) of insertions for these TdT variants revealed a wide spectrum of nucleotide preferences and average insertion lengths (Figure 1c). A/T biases (assuming A/T content in insertions ≅ A/T bias of the TdT variant) ranged from 23% to 87%, and the average lengths of insertions varied from 1-5 bp (Figure 1c). We provide all performance metrics for all unique TdT variants we analyzed in Supplementary Tables 1-3. The results of this experiment yielded multiple A/T-biased, G/C-biased, and unbiased TdT variants for application (see below).

Given the high-throughput nature of our assay, we wished to examine the patterns of TdT mutations in our libraries responsible for altering biases. To do so, we grouped TdT variants from each library into 3 different bias bins representing G/C-biased, A/T-biased, and unbiased insertion sequence outcomes (Figure 1c,d). Specifically, the ∼100 most G/C-biased, A/T-biased, or unbiased variants for each library were grouped into their respective bins (some bins contained up to 105 library members, instead of exactly 100, because the biases of several variants at the boundary of a given bin were identical). Library 2 had only 26 variants with a G/C bias (<50% A/T), so 26 instead of 100 variants are in this bin. The composition of amino acids at each randomized position for each bin was calculated to determine if certain mutations are over- or under-represented in different bias profiles. The resulting heatmaps of amino acid compositions (Figure 1d) showed that D399 is conserved among G/C-biased TdTs, suggesting it plays a role in effecting the natural G/C bias of TdT. This is further supported by the fact that D399 is usually mutated in the A/T-bias bins. In library 3, all 3 mutagenized positions (R454, E457, R461) frequently maintained their wild-type identities in the G/C-bias bin but were depleted in the A/T-bias bin, suggesting that all three residues, which point toward the nucleobase of the incoming nucleotide during polymerization, have an effect on maintaining the native G/C bias of TdT and act as hotspots for changing this bias to A/T.

### Properties of TdT variants determined in the multiplex assay are reproduced when assayed individually

To validate the results of the multiplex assay, TdT variants of interest were individually constructed and cloned into a mammalian expression vector. Overall, 74 library members were selected for individual characterization. They were chosen based on their A/T biases and average insertion lengths, aiming for a diverse range in both criteria, while also prioritizing for high activity and number of insertions sequenced (Supplementary Table 4). Each TdT variant of interest was then co-transfected into HEK293T cells alongside a construct encoding Cas9 and a sgRNA targeting the genomic HEK3 site. Cell populations were collected 3 days later and editing outcomes at the HEK3 site were assessed *via* NGS. We observed a strong correlation between the variants’ performance in the multiplex assay and the individual assays here for A/T bias (Pearson correlation coefficient, ρ = 0.952) and insertion length (ρ = 0.915) (Figures 2a and 2b). For A/T bias, the regression coefficient was 0.878 (95% confidence interval = 0.812-0.945) while for insertion length, the regression coefficient was 0.576 (95% confidence interval = 0.516-0.635), reflecting the observation that insertion lengths were shorter in the individual assays than in the multiplex assay (Figure 2b). This is likely due to the different methods for expressing TdT used in the multiplex assay (lentivirus) versus the individual assays (transient transfection). Because TdT is thought to use a distributive mechanism for polymerization^2^, if lentiviral expression in the multiplex assay achieved higher intracellular concentrations of TdT, the over-expression could promote longer insertions. Nonetheless, the A/T bias and insertion length measurements from the multiplex assay can predict those of the individual assays, validating the multiplex assay as a way to isolate valuable TdT variants for application.

**Figure 2.**
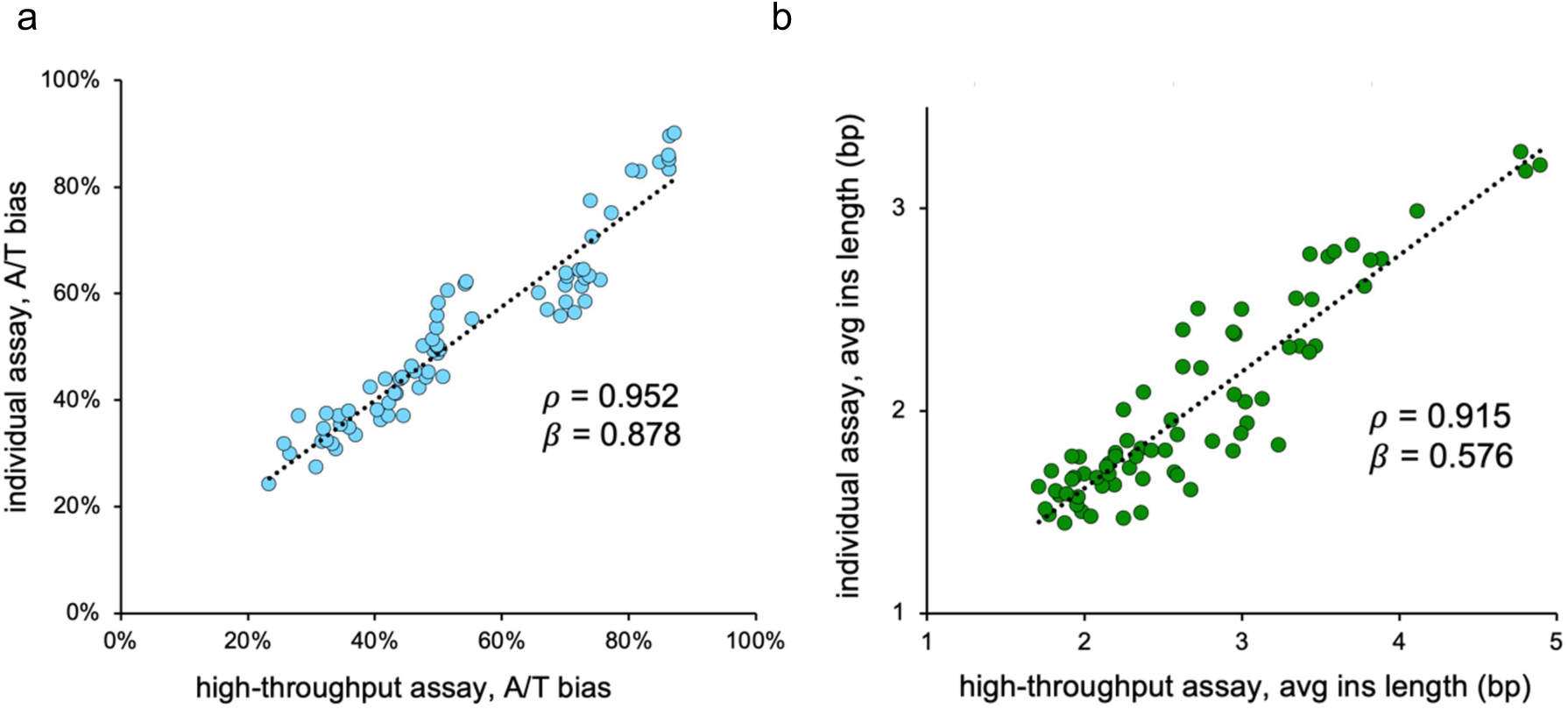
Comparison of mutant TdT properties between the high-throughput multiplex assay and individual assays, **a,** 74 TdT variants were individually assayed in a follow-up to the high-throughput screen, and the apparent A/T biases of these variants (% of inserted nucleotides comprised of A/T) are plotted for both assay methods, **b,** The average insertion length for each of the 74 individually-assayed variants is shown for both assay methods, *ρ,* Pearson’s correlation coefficient. *β*, regression coefficient.

### *In vitro* biases of TdT mutants match those determined in the *in vivo* multiplex assay

When expressed in *E. coli*, purified, and assayed biochemically, the TdT variants recapitulated their A/T biases determined in the multiplex screen. We chose 7 TdT mutants to test *in vitro*, plus wild-type TdT for comparison. A TdT variant with nucleotide preferences nearly identical to wild-type was included to investigate if variants with similar performance *in vivo* also appear similar *in vitro*. All other TdT variants were selected because they had extreme biases or were unbiased. Two of the most A/T-biased variants from the pooled screen – A/T-biased1, with R454L, E457K, R461L and 87% A/T bias; and A/T-biased2, with R454I, E457K, R461A and 86% A/T bias – were included. Since dATP is the most disfavored nucleotide for wild-type TdT^2^, we reasoned that these A/T-biased variants may be broadly useful for boosting the efficiency of *in vitro* DNA synthesis by TdT. Also, 3 unbiased variants – Unbiased1, with R454M, E457C and 50% A/T bias; Unbiased2, with D396R, L398F and 50% A/T bias; and Unbiased3, with R454K, E457R and 50% A/T bias – were selected for the *in vitro* assay because equal incorporation of all bases would increase the entropy of TdT-generated sequences. We envisioned these could aid in synthesis of aptamer libraries^5^. In addition, G/C-biased variants are desirable subjects for *in vitro* testing, since G-biased TdTs may improve the performance of biosensors that synthesize G-quadruplexes for signal generation^16,46–57^ or enable the use of these biosensors in crude biosamples (without needing to purify specimens to remove all 4 natural nucleotides). However, because G/C-biased variants were rare among the library members we profiled in our multiplex assay, we attempted to amplify G/C bias by creating a double mutant possessing mutations from two modestly G/C-biased variants isolated from libraries 2 and 3 (L398M from library 2, plus R454D and E457D from library 3). We ultimately obtained one double mutant – G/C-biased1, with L398M, R454D, E457D – that was marginally more G/C-biased than its parent mutants, with an A/T fraction of 28% when tested individually in HEK293Ts (Supplementary Figure 3). We used this double mutant for subsequent experiments.

For all TdT variants, the overall biases were faithfully reproduced when assayed *in vitro*. There was a strong correlation between the A/T biases observed *in vitro* and *in vivo* (Pearson’s ρ = 0.950), and the regression coefficient was 0.841 (95% confidence interval 0.564-1.118) (Figure 3a). We note that, because the *in vitro* assay profiled TdT’s usage of all 4 nucleotides on a ssDNA primer instead of at a DSB, we gained additional information about how TdT variants prefer A vs. T and G vs. C. When TdT acts on a DSB, as in the *in vivo* multiplex screen, the presence of 2 available 3’ DNA ends at the DSB obscures the exact identity of TdT-mediated insertions (Supplementary Figure 4). Specifically, TdT’s preferences between complementary bases cannot be discerned, since an inserted base on the top strand of DNA at the DSB would appear identical to insertion of its complementary base on the bottom strand. For this reason, we recommend *in vitro* assaying for applications where the ranking among all 4 bases must be precisely known.

**Figure 3.**
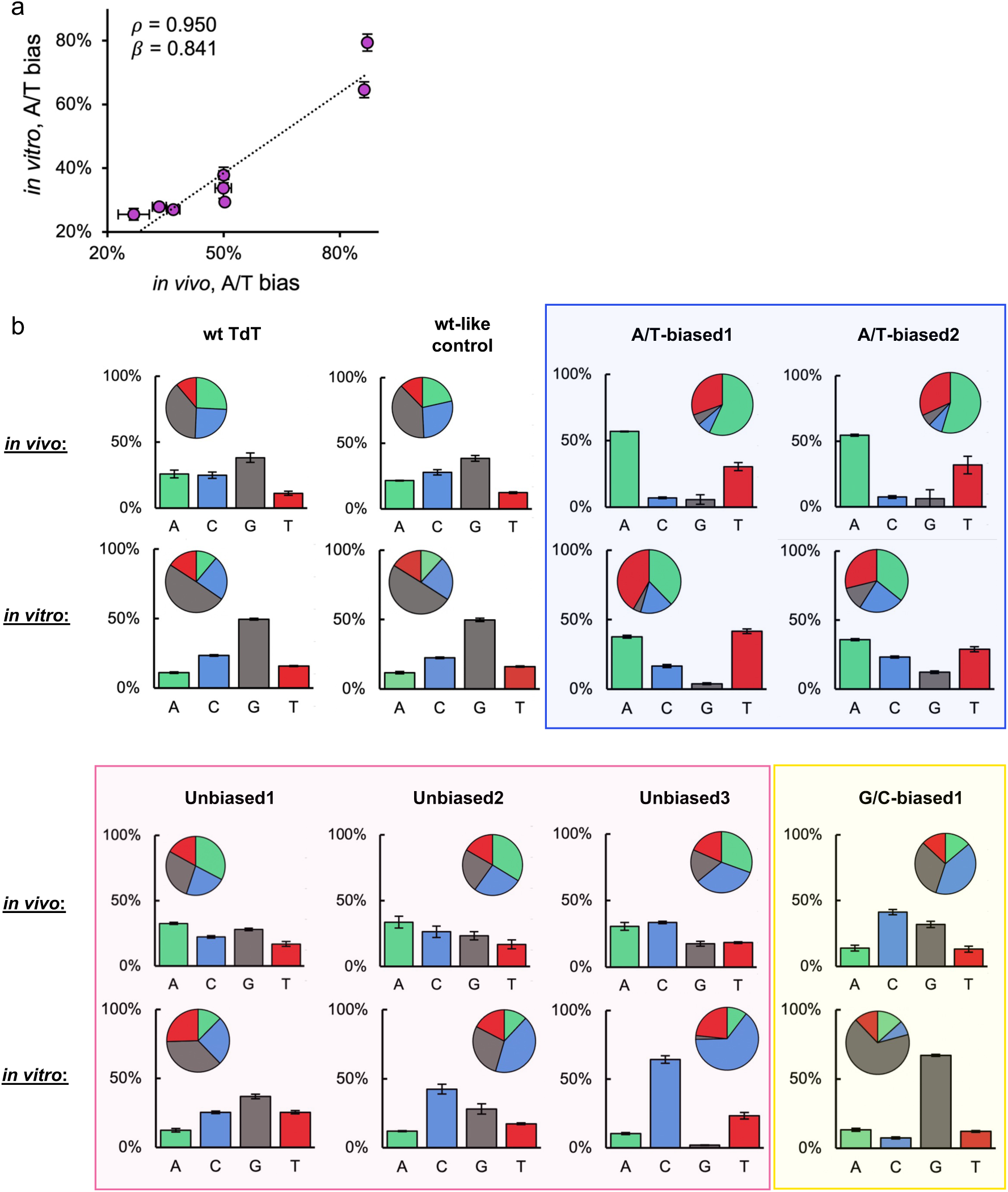
*In vitro* assays of TdT mutants with practically useful nucleotide preferences (*i.e*., having strong biases or equal preferences for A/T and G/C). **a,** Relationship between A/T biases observed *in vivo* and *in vitro, ρ,* Pearson’s correlation coefficient, *β,* regression coefficient, **b,** Bar and pie graphs both depict the percent of incorporated nucleotides comprised of A, C, G, or T for each variant. Except for the G/C-biased variant, *in vivo* data is from the massively parallel pooled screen in HEK293T cells. The G/C-biased variant is a double mutant (combining mutations from variants taken from two libraries), so the *in vivo* results are necessarily from an individual assay (3 biological replicates). Error bars indicate the standard deviation for 3 technical replicates of the high-throughput *in vivo* assay, and 3 biological replicates for the *in vitro* assay.

### Engineered TdTs improve barcode diversity for DNA recording in live cells

Cell HistorY Recording by Ordered iNsertion (CHYRON) is a DNA recorder that our group previously established (Figure 4a)^36^. CHYRON works through the iterative accumulation of random insertion mutations, generated by TdT, at a neutral locus in the genome. These insertions can be used to infer lineage relationships among cells. Iterative accumulation of insertions is possible because the recording locus encodes a homing guide RNA (hgRNA) that, once edited by Cas9 and TdT, becomes a mutated hgRNA that can direct further editing at the same locus. The original version of CHYRON used wild-type TdT for insertion, but as expected, we observed a clear G/C bias in the growing insertions at the CHYRON locus. This bias increases the probability that unrelated cells generate identical barcode sequences, resulting in the false appearance of shared ancestry (homoplasy). We hypothesized that our newly discovered unbiased TdT variants could be used in the CHYRON framework to generate more random insertion sequences with higher entropy.

**Figure 4.**
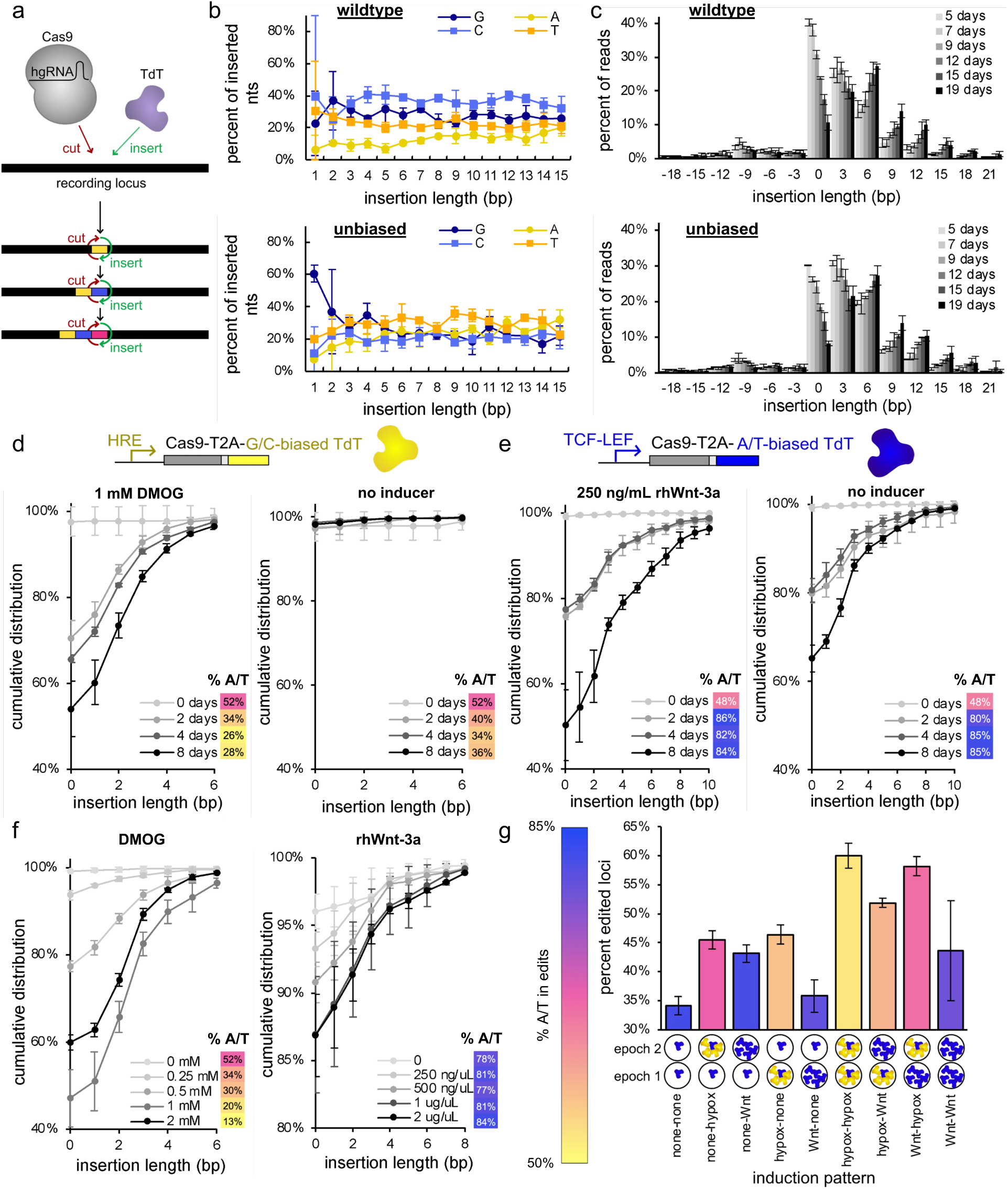
Altered-bias TdTs were used for DNA recording in live HEK293Ts. **a,** Depiction of CHYRON, an *in vivo* DNA recorder that iteratively acquires insertion mutations through the joint activity of Cas9, a homing guide RNA (hgRNA), and TdT. **b,** Proportions of inserted bases for two different versions of CHYRON– the legacy CHYRON that uses wildtype TdT, and a new version that uses an unbiased variant of TdT (Unbiased3). After 19 days of continuous mutation accumulation with these two systems, the compositions of insertions were assessed. An unbiased TdT was tested because the random insertions created by CHYRON can be used to record cell lineage relationships, and an unbiased composition of inserted nucleotides would improve information content. A perfectly unbiased TdT would produce insertions with 25% each base across all insertion lengths. **c,** Distribution of insertion lengths over time when CHYRON was propagated with wild-type TdT or the unbiased variant of TdT, as in b. **d,** A G/C-biased TdT (G/C-biased1) and a hypoxia-inducible promoter (4xHRE_YB TATA) were installed into the CHYRON architecture, and various durations of induction with a chemical hypoxia mimic (DMOG) were recorded. **e,** An A/T-biased TdT (A/T-biased2) and a Wnt-inducible promoter (7xTCF/LEF) were used to record various durations of exposure to recombinant Wnt protein (rhWnt-3a). **f,** Different concentrations of DMOG and Wnt were recorded in an analog fashion, using the appropriate inducible CHYRON constructs with biased TdTs. **g,** Both hypoxia- and Wnt-inducible CHYRON cassettes with differently-biased TdTs were transfected into a single population of HEK293Ts, and different patterns of induction were recorded. The cells were exposed to hypoxia, Wnt, or no inducer for 4 days (1 “epoch”), then they were re-transfected with the CHYRON cassettes and again exposed to hypoxia, Wnt, or nothing for an additional 4-day epoch. The extent of editing, and the A/T content of the insertion sequences, are graphed. Logos beneath the x-axis show the expected profile of expressed TdTs in each epoch for each pattern of induction. Blue = Wnt-inducible and A/T-biased TdT, yellow = hypoxia-inducible and G/C-biased TdT. The Wnt-inducible promoter is leaky, so basal expression of the A/T-biased TdT is always expected. Panels b-f show the averages of 3 technical replicates, and panel g shows the averages of 5 technical replicates. Error bars indicate the standard deviation among replicates.

TdT variants Unbiased1, Unbiased2, and Unbiased 3 (Figure 3) were tested in the CHYRON context through the establishment of stable cell lines containing a single hgRNA (the CHYRON recording locus) encoded at AAVS1 and pCAG-Cas9-T2A-TdT cassettes integrated by piggyBAC transposition. The CHYRON hgRNA started with a 20-nt spacer, leading to high overall activity and accumulating an average of 5 inserted nucleotides^36^. These cell lines were continuously passaged for 19 days. Periodically, subpopulations were sequenced at their CHYRON loci. All of the unbiased TdT variants increased the per nucleotide diversity of CHYRON loci relative to the wild-type TdT at the end of the 19-day experiment. Wild-type TdT’s insertions reflected an overall A/T bias of 37%, and a corresponding Shannon entropy of 1.93 bits per inserted nucleotide (Figure 4b). On the other hand, the best performing unbiased TdT (Unbiased3) achieved 1.99 bits per inserted nucleotide (compared to 2 bits for the theoretical maximum). Since Unbiased3 was also found to grow the CHYRON locus with similar efficiency as wild-type TdT (Figure 4c), its unbiased nature represents an improvement over the legacy CHYRON system. Although the difference between 1.93 bits (wild-type TdT) and 1.99 bits (Unbiased3) per inserted nucleotide is small, we note that insertions contain multiple nucleotides and in systems such as DARLIN^3^, multiple loci are used for TdT insertions. For example, CHYRON loci in this experiment frequently reached 6 bp in length (Figure 4c) for which any given observed sequence should be considered one among ∼3,000 at 1.93 bits per nucleotide (11.58 bits per 6-nt insertion = 2^11.58^ sequences) versus one among ∼4,000 sequences for 1.99 bits per nucleotide (11.94 bits per 6-nt insertion = 2^11.94^ sequences). In other words, the small difference in Shannon entropy per inserted nucleotide compounds exponentially in information content of actual insertion outcomes.

### TdT mutants with extreme nucleotide insertion preferences enable dual-channel signal recording in HEK293T cells

In previous work, we used our *in vivo* DNA recorder, CHYRON (Figure 4a), as an analog recorder for hypoxia^36^. We demonstrated that the combined activities of a hypoxia-inducible Cas9, a hypoxia-inducible TdT, and a constitutively expressed hgRNA produce a hypoxia dose- and duration-dependent increase in the number and size of insertions at recording loci. The hypoxia-recording experiments served as a first proof of principle for analog signal recording, but in theory, CHYRON could be used to record any event that causes a transcriptional response in the cells of interest. Provided that the corresponding transcription factor binding sites are known, recording the desired signal only requires substituting the appropriate inducible promoter into the Cas9 and TdT expression cassette. Thus, CHYRON has the potential to record many different signals. However, with one TdT, one can only record one signal at a time (or the sum of all individual signals). We aimed to alter this paradigm by linking different signals to the expression of differently biased TdTs.

We used an A/T-biased TdT variant, specifically A/T-biased2 (86% A/T), to record Wnt, and a G/C-biased TdT variant, specifically G/C-biased1 (28% A/T), to record hypoxia. The variants were cloned into inducible cassettes with the general architecture [inducible promoter]-Cas9-T2A-[TdT variant]. With this setup, we envisioned that the amount and identities of insertions could be used to decipher the cells’ histories of induction by two signals.

Before testing the biased TdTs’ ability to perform dual-signal recording when jointly implemented, we first assessed their individual performance as CHYRON analog recorders in separate populations of cells. The recording cassettes were transiently transfected into HEK293Ts that already possessed a CHYRON locus – a constitutively expressed hgRNA gene stably integrated in the genome^36,58,59^– to enable iterative rounds of editing. These populations of cells were either exposed to the relevant inducer or passaged in inducer-free media and sequenced *via* NGS at various timepoints over the course of 8 days. Encouragingly, when the hypoxia-inducible cassette with the G/C-biased TdT was induced with DMOG, the cumulative distribution of insertion lengths shifted towards more and longer insertions as induction duration increased, and very little insertion was observed without DMOG (Figure 4d). The A/T content of the insertions at the CHYRON loci was consistently low over time, staying between 26% and 34% A/T after 2+ days of induction, in agreement with the expectation that insertions are generated by the G/C-biased TdT. We note that the small amount of short insertions observed at day zero of DMOG induction or in the uninduced cases were not as GC-biased as the longer insertions. This is because when TdT misses an opportunity to edit a DSB, non-homologous end joining (NHEJ) often results in short insertions with ∼50% A/T content, skewing the bias away from that of the specialized TdT. Overall, the behavior of the hypoxia-recording cassette allowed different durations of DMOG exposure to be classified from the CHYRON records, and the presence or absence of inducer was clear based on the extent of editing.

When the same experiment was conducted for the Wnt-inducible cassette with an A/T-biased TdT (Figure 4e), the resolution of recording was somewhat coarse, as 2 days of induction was indistinguishable from 4 days of induction by the cumulative distributions of insertion lengths. However, 0, 2 or 4, and 8 days produced distinct compositions of insertion lengths, so we consider the resolution of this recorder to be approximately 4 days. The A/T content of the insertions was consistent with the expected values for the A/T-biased TdT variant (82-86% at 2+ days of induction). Unfortunately, when an identical population of cells was passaged in the absence of inducer, a considerable amount of leaky editing occurred. This complicates the interpretation of recorded insertions, but it doesn’t preclude the use of this Wnt recorder, as editing still proceeded at a slower pace in the absence of inducer – for each timepoint, the induced population was more extensively edited than the uninduced population (Figure 4e). Future work will focus on reducing the leakiness of the Wnt promoter.

Next, we tested the two distinct recorders, again in separate populations, but this time for their responsiveness to different doses of inducer (Figure 4f). Populations of HEK293Ts were transfected with an inducible cassette, induced with a range of inducer concentrations, then collected and sequenced *via* NGS after 2.5 days of induction. The dose responses of both recorders showed increased editing (both overall percent edited loci and length of insertions) in the presence of higher inducer concentrations. There was one exception to this trend – the hypoxia recorder produced less editing with 2 mM DMOG than with 1 mM DMOG in the culture media. We believe this was due to the cytotoxic effects of DMOG on cells, as we observed high rates of cell death in the cultures with high DMOG concentrations. Insertions in the hypoxia recording experiment trended towards decreased A/T content in response to increased inducer (34% A/T for 0.25 mM DMOG, compared to 13% A/T for 2 mM DMOG), consistent with the greater action of the G/C-biased TdT when highly induced against the natural ∼50% A/T NHEJ short insertion outcomes when TdT does not engage. In the case of the Wnt recorder, the A/T content of the insertions is similar across all induction levels (77-84% A/T), suggesting that the action of the A/T-biased TdT is enough to overcome the ∼50% A/T NHEJ short insertion outcomes even when not induced (due to the leakiness of the Wnt promoter used).

Having confirmed that the hypoxia recording cassette can generate G/C-rich insertions and the Wnt recording cassette can generate A/T-rich insertions in response to their corresponding signals, we next performed dual-signal recording of both hypoxia and Wnt in a single population of HEK293Ts. For this, populations of cells were co-transfected with both recording cassettes, then induced with hypoxia, Wnt, or nothing for 4 days (1 “epoch”). Then, the populations were re-transfected with the recording cassettes and subjected to an additional epoch of induction with hypoxia, Wnt or nothing. The CHYRON loci in the 9 final populations were sequenced and inspected (Figure 4g). The percent of CHYRON loci bearing an edit generally correlated with the amount of time the populations were induced by either signal. For 8/9 populations, the total percent of CHYRON loci with an edit followed the trend 0 epochs < 1 epoch < 2 epochs of induction. In addition, the populations recorded the identities of the inducers in each epoch, as reflected in the A/T content of the CHYRON records. For example, every sample that exclusively experienced Wnt induction (none-Wnt, Wnt-none, and Wnt-Wnt) had insertions comprised of 81-83% A/T. On the other hand, every sample that exclusively experienced hypoxia (none-hypox, hypox-none, and hypox-hypox) produced a lower amount of A/T rich insertions, with the hypox-hypox conditions producing the lowest (54% A/T). We note that, because of the leakiness of the Wnt promoter (Figure 4e-f), the conditions that experienced only hypoxia did not reach an A/T content of 28% even though that is the expected A/T content of insertions by the G/C-biased TdT used to record hypoxia. Still, this experiment showed that we can distinguish between whether a cell saw hypoxia or Wnt if only one was present.

For samples exposed to both hypoxia and Wnt (hypox-Wnt and Wnt-hypox), CHYRON records had intermediate A/T content (61-72% A/T). Furthermore, samples that experienced no induction (none-none) could be identified by the overall low percent of editing, but given the leakiness of the Wnt promoter (Figure 4e-f), the A/T content (83%) of these samples was not informative on the lack of induction.

While inspecting the A/T biases of the records, we also noticed that the populations exposed to hypoxia for only 1 epoch had a lower A/T content if hypoxia was the first epoch (hypox-none, 59% A/T) compared to when it was the second epoch (none-hypox, 74% A/T). This is likely because growth of the CHYRON locus through sequential editing slows over time^36^ such that early TdT action had an outsized influence. Because of this slowdown, we could not confidently reconstruct the order of inducers by looking at how the A/T content changes across the length of individual insertions generated through multiple insertion rounds. Future work on engineering CHYRON to maintain a constant rate of editing and reducing leaky recording of Wnt should lead to better recording performance. Overall, these data show that we can fully or partially decipher the signals (hypoxia and/or Wnt) our populations of cells experienced by characterizing the length and A/T content of CHYRON loci, while highlighting areas for improvement in this approach.

## Discussion

We have described a new approach to screening libraries of TdT variants for changes in activity and bias with unprecedentedly high throughput. Using this screening method, we successfully identified TdTs with advantageous properties for our application of interest, *in vivo* DNA recording with CHYRON^36^, and simultaneously generated a database describing the activities, insertion biases, and average insertion lengths of 25,623 TdT variants. Although our multiplexed assay takes place in HEK293T cells, we demonstrated that the insertion biases *in vivo* were also reflected *in vitro*. The TdT variants we have already characterized, and the multiplexed assay we developed, should both aid in the development of other *in vivo* and *in vitro* applications of TdT.

Among the TdT variants screened in this study, we captured a wide variety of nucleotide preferences (propensity to insert A vs. T vs. C vs. G) and average insertion lengths. Besides the CHYRON DNA recording applications shown, we imagine extensions of our new TdTs to other technologies. For example, our altered-bias variants can be used for dual signal recording in the DARLIN architecture^3^, which uses Cas9 plus TdT to generate random insertions at many recording sites across the genome. Additionally, our A/T-biased TdTs could potentially be substituted into enzymatic oligo synthesis platforms to overcome the currently slow rate of dATP addition (dATP requires the longest coupling time out of all 4 natural nucleotides, 2x longer than the others)^60^. Finally, our multiplex assay could be used to profile the activities and biases of naturally occurring TdT variants, which may have profound impacts on T- and B-cell receptor diversity (and therefore, adaptive immune function) among humans. To date, only a handful of such TdT variants have been characterized, and only *in vitro*^61^.

## Methods

### Clonal plasmid construction

Enzymes for cloning were procured from NEB. Constructs were assembled by PCR-amplifying the fragments with primers encoding SNPs or larger mismatches (*e.g.*, spacer sequences in sgRNAs were encoded as 20-bp primer overhangs on the ends of amplicons), then assembling fragments *via* Gibson Assembly or Golden Gate Assembly. After confirming faithful assembly *via* Sanger sequencing (Azenta Life Sciences) or whole-plasmid sequencing (Primordium Labs), plasmids were hp-miniprepped (GenElute HP Plasmid Miniprep Kit, NA0150) to produce high-purity plasmid stocks for downstream transfections.

The only sgRNA needed in this study was one targeting the HEK3 sequence (Supplementary Table 5); all *in vivo* experiments targeted Cas9 to either the endogenous HEK3 locus, or a copy of the HEK3 sequence inserted into a synthetic context. The sgRNA was cloned *via* Gibson Assembly, adding the appropriate spacer sequence into the pSQT1313 vector (Addgene #53370)^62^, which was a gift from Keith Joung.

For *in vivo* assays of TdT variants with Cas9 targeting the HEK3 locus, we used one of our previously established plasmids that harbors the wild-type hTdT gene in a pcDNA3.1 backbone (Addgene #126450)^36^. Point mutations were introduced into this construct to constitute variants of interest.

For TdT variants that were installed into the CHYRON architecture, a piggyBAC expression vector from the Mammalian Toolkit^63^ was used. The Mammalian Toolkit was a gift from Hana El-Samad (Addgene kit #1000000180). For continuous propagation of cells containing the CHYRON locus, the pCAG promoter was used (Addgene #123700, also a Mammalian Toolkit part plasmid) to drive expression of a Cas9-T2A-TdT cassette, which was PCR-amplified from one of our previously described plasmids (Addgene #132667)^36^. Golden Gate Assembly was used to first make the piggyBAC-pCAG-Cas9-T2A-TdT plasmid with wild-type hTdT, which was then used as a template for generating CHYRON cassettes encoding variants of interest. For dual signal recording, the pCAG promoter was replaced with 4xHRE_YB TATA for hypoxia recording (Addgene #117399)^64^ or 7xTCF/LEF for Wnt recording (Addgene #12456)^65^. M50 Super 8x TOPFlash containing the 7xTCF/LEF promoter was a gift from Randall Moon, and 4xHRE_YB TATA-sfGFP-CMV_dsRed was a gift from Yvonne Chen. Also, an oxygen-dependent degron from one of our previously described plasmids (Addgene #132667)^36^ was appended to Cas9 in the hypoxia-recording cassette. All pertinent plasmids from this study will be made available on Addgene.

For *in vitro* assays, a truncated version of the gene for wild-type human TdT was codon-optimized for expression in *E. coli* B (Integrated DNA Technologies, https://www.idtdna.com/CodonOpt) and synthesized as a gBlock from Integrated DNA Technologies (IDT). The N terminus was excluded, as we have found the BRCT domain^66^ compromises expression, and it is unnecessary for observing TdT activity *in vitro*. In the final design, only residues 125-509 were included. The TdT gene was cloned into a pET28a expression vector (Addgene #69864-3) with an N-terminal 10xHis tag, linker, and protease cleavage site. The wild-type TdT plasmid was used as a template to produce plasmids for each variant of interest.

### Site saturation mutagenesis library cloning and lentivirus production

All cloning enzymes and buffers were procured from New England Biolabs. To construct a template for subsequent library cloning, a wild-type human TdT (hTdT) lentiviral expression plasmid was assembled. The hTdT sequence was amplified from one of our previously reported plasmids (Addgene #163643)^36^. The pCMV Blast Dest lentiviral expression vector (Addgene #17451)^67^, which was a gift from Eric Campeau and Paul Kaufman, was used as the starting point for the backbone. Several modifications to the hTdT gene and expression vector were introduced *via* PCR with point mutations encoded in the primers, then the amplicons were stitched together in a Gibson Assembly. Specifically, new SalI and PstI restriction sites were added (using synonymous point mutations), to enable downstream cloning steps, while endogenous SalI and PstI restriction sites were removed. Also, the ampicillin resistance gene in the lentiviral backbone was replaced with kanamycin resistance, an sfGFP gene was added for virus titering, and a puromycin resistance gene was added for later selection of transduced cells (Supplementary Table 5).

To achieve site saturation mutagenesis, we PCR-amplified the wild-type hTdT gene in two consecutive PCRs, using primers that introduced the degenerate bases NNK at the codons we wished to randomize (Supplementary Table 6). N represents any nucleotide, and K represents G or T. The last nucleotide in the randomized codons is designated with K rather than N because this allows all 20 natural amino acids to be included, while reducing library size and avoiding most stop codons^68^. We chose to construct the library in two sequential PCRs to minimize bias in the NNK sequences. In the first PCR, the TdT gene was amplified from its promoter to the last nucleotide upstream of the NNK codons. This PCR product was digested with DpnI to remove wild-type template plasmid, then purified *via* extraction from an agarose gel (Epoch Life Sciences All-In-One DNA Only Spin Columns). The purified amplicons were used as the template for a second PCR that introduced NNK sequences as primer overhangs, thereby avoiding biases that could arise due to differences in melting temperatures among library members that could base pair with the wild-type template. These primers also introduced restriction sites for the downstream plasmid assembly. Primers were synthesized by Integrated DNA Technologies (IDT), with hand-mixed degenerate bases harboring a 1:1:1:1 composition of A:C:G:T for N and 0:0:1:1 for K sites in the NNKs. The PCR products were purified on Epoch Life Sciences All-In-One DNA Only Spin Columns.

Two different approaches were implemented to assemble the mutagenized TdT genes into the lentiviral expression backbone. Libraries 1 and 2 were cloned *via* restriction digestion with SalI-HF and PstI-HF followed by ligation with T4 DNA Ligase. For this, the inserts and backbones were digested in separate reactions with 1 µg of DNA per 50 uL reaction, 100 U of SalI-HF, 20 U of PstI-HF, and Cutsmart buffer. For the plasmid backbones, a miniprep of the wild-type TdT plasmid was digested directly, and calf intestinal alkaline phosphatase (CIP) was added to the digestion to prevent downstream self-ligation of the empty vector. The digestion was incubated at 37°C for 1 hour. Digestion products were extracted from agarose gels, then the inserts and backbones were ligated together with 220 ng of backbone, 110 ng of insert, 1760 U T4 Ligase, and T4 ligase buffer in 44 uL reactions, for a total of 4 ligation reactions per library. In addition, a vector-only ligation was set up for each library with the same conditions, but no insert (to measure the rate of empty vectors self-ligating). The ligation was carried out at 4°C overnight, followed by incubation at 65°C for 10 minutes. For library 3, plasmids were constructed *via* Golden Gate Assembly because synonymous SalI and PstI sites were inaccessible near the NNK target sites. The backbone for library 3 was amplified *via* PCR while simultaneously adding BsmBI restriction sites as primer overhangs, then it was digested with DpnI and gel extracted.

Golden Gate Assembly was conducted in 20 uL reactions with 26 ng vector DNA, 12 ng insert DNA, T4 DNA ligase buffer, 10 U BsmBI, and 400 U T4 DNA Ligase, for a total of 32 reactions. A vector-only Golden Gate Assembly was also tested. Thermal cycling conditions were (42°C for 5 min, 16°C for 5 min) x30 cycles, 55°C for 5 min, and 80°C for 10 min.

For all three libraries, the assembled plasmids and empty vector controls were cleaned and concentrated *via* ethanol precipitation with yeast tRNA (ThermoFisher) as a co-precipitate (12 µg yeast tRNA per 300 ng assembled DNA per 840 uL precipitation). The final DNA + tRNA was resuspended in 4.1 uL ddH2O. For 4 transformations per library, 1 uL of this concentrated DNA + tRNA was transformed into 20 uL of ElectroMAX Stbl4 cells according to manufacturer’s instructions (ThermoFisher). The transformation outgrowth was incubated for 1 hour at 30°C. After the outgrowth, all transformations for each library were pooled together, then plated on 8 large LB agar + kanamycin dishes (15 cm diameter) per library. Serial dilutions of the transformations were also plated on LB agar + kanamycin to allow colony counting and library coverage calculations. Plates were incubated at 30°C for approximately 24 hours. All colonies from the large plates were scraped and pooled together in 4 mL 2YT + kanamycin per plate. These cell suspensions were midiprepped (GenElute HP Plasmid Midiprep Kit, Millipore Sigma), and the final pools of library DNA were used for lentivirus production and titering at the University of San Francisco Viracore. Immediately upon receiving the virus, it was divided into single-use aliquots (to avoid repeated freeze-thaw cycles), then stored at -80°C.

### High-throughput pooled screen of TdT libraries in HEK293Ts

HEK293Ts were procured directly from ATCC (CRL-3216) but were not otherwise authenticated. All cell culture was conducted in DMEM with high glucose plus GlutaMAX (Gibco, 10566024) and 10% FBS (Sigma, 12306C). Cultures were maintained at 37 °C and in 5% CO_2_.

At the start of the high-throughput screen, cryopreserved HEK293Ts were placed in a 37 °C bead bath until nearly thawed, then resuspended in 10 mL pre-warmed DMEM with high glucose and GlutaMAX Supplement (Gibco, 10566024), plus 10% FBS (Sigma, 12306C) and seeded into a cell culture treated T-75 flask (Corning, 430641U). The initial population of 293Ts was grown long enough to passage once before transducing with lentivirus (5 days). Transduction was carried out in 3 T-150 flasks per library (Corning, 430825). When the cultures were approximately 30% confluent, the lentiviral TdT libraries were transduced at a multiplicity of infection (MOI) of 0.3, with 8 µg/mL polybrene (Fisher Scientific, TR1003G) supplemented to the media. 24 hours post-transduction, 1/3 of the cells in each T-150 were passaged into a new T-150 (a stringent dilution was avoided, to avoid losing library members). At 48 hours post-transduction, the media was replaced with fresh culture media containing 2 µg/mL puromycin (InvivoGen, ant-pr-1). After approximately 1.5 days of selection, dead cells were removed by refreshing the puromycin-containing media. Cultures were maintained in 3 T-150 flasks per library, passaging as needed for a total of 6 days in the presence of puromycin. For transfection with Cas9 and an sgRNA designed to generate DSBs downstream of the TdT genes, each library was passaged for the final time into 3 T-150 flasks, aiming for 10% seeding density. Approximately 24 hours later, 11.1 µg of a Cas9-expressing plasmid (Addgene #41815)^69^, 7.4 µg of the appropriate sgRNA-expressing plasmid (Supplementary Table 5), and 1 µg of an mCherry-expressing plasmid (included for a visual indicator of the transfection efficiency) were transfected into each flask with FuGENE HD transfection reagent (Promega E2312), according to the manufacturer’s instructions, with a 15-minute complexing time. The Cas9-expressing plasmid was a gift from George Church. 16 hours after transfecting, the culture media in the flasks was refreshed. 3 days post-transfection, the cultures were trypsinized, resuspended in regular culture media, and a small fraction (4% of the cells) was seeded into a T-25 flask (Thermo Scientific, 156340) for Mycoplasma testing (Applied Biological Materials, C214). The remainder of the cells were washed once with phosphate buffered saline (ThermoFisher, 10010023), then the cell pellets were frozen at -80°C for later genomic DNA extraction and sequencing. All Mycoplasma tests were negative.

### Cell culture methods for individual TdT assays and CHYRON experiments

To assay individual TdT variants, cryopreserved HEK293Ts were placed in a bead bath at 37 °C until nearly completely thawed. Cells were gently resuspended in pre-warmed culture media and seeded into the appropriate size cell culture treated dish to achieve a seeding density of ∼50%. 1-2 days later, cultures were split into 6-well dishes (Falcon, 353046) with a seeding density of approximately 5%. The following day, hp-minipreps of Cas9 (750 ng), the HEK3-targeting sgRNA (500 ng), and the TdT variant of interest (750 ng) were transiently transfected using FuGENE HD Transfection Reagent (Promega E2312) according to manufacturer instructions. 20-minute complexing times were used. 3 days post-transfection, the cells were collected by aspirating the media out of the cultures, spraying cells off the plates with cold phosphate buffered saline (ThermoFisher, 10010023), pelleting the cells, removing the supernatant, and placing cell pellets at -80°C for later analysis.

For CHYRON experiments, we used our previously reported CHYRON_20i_ cell line^36^, which possesses a homing guide RNA (hgRNA) gene^58,59^ with a 20 nt spacer integrated at the AAVS1 locus and chromatin insulators^70^. The Cas9-T2A-TdT transgene(s) with the TdT variant(s) of interest were either stably integrated into the genome with Super PiggyBAC Transposase (System Biosciences Innovation, PB210PA-1) (for long-term propagation of the CHYRON locus as in Figure 4b-c) or transiently transfected at 4-day intervals (for signal recording as in Figure 4d-g). In both cases, FuGENE HD Transfection Reagent with a 20-minute complexing time was used to introduce the transgenes into the cells (Promega E2312). For stable cell lines, samples were passaged into tissue culture treated 24-well dishes (Fisher Scientific FB012929) 1 day prior to transfection, with a seeding density of approximately 20%. Transfection was performed with a 5:1 cargo: transposase ratio. 2 days later, samples were passaged into 6-well dishes, with media containing 10 µg/mL blasticin (Thermo Scientific, J672168EQ), to select for successfully transformed cells. An untransfected control was also cultured in 10 mg/mL blasticin to confirm the selection was carried out for a long enough duration to kill all non-resistant cells in the experimental samples before the first timepoint was collected. Blasticidin selection was maintained throughout the entire timecourse. Cultures were passaged approximately once every 3 days, and cell pellets containing at least 300,000 cells (corresponding to 10% confluency in a 6-well dish) were periodically sampled for NGS. For transiently transfected samples (signal recording experiment), the CHYRON_20i_ cells^36^ were passaged into 24-well dishes 1 day prior to transfecting. FuGENE HD transfection reagent (Promega E2312) was used as specified in the user manual to transfect 400 ng of the hypoxia and/or Wnt recording cassettes into the 24-well cultures, with a 20 min complexing time. 16 hours after transfecting, the culture media was aspirated off and replaced with either regular culture media or inducer-containing media. Recombinant human Wnt-3a protein (R&D Systems, 5036-WN-010) was used to induce Wnt recording at a final concentration of 250 ng/mL in the culture media, unless stated otherwise. The HIF-hydroxylase inhibitor, DMOG (Millipore Sigma, 40009150MG), was used to induce hypoxia recording at a final concentration of 1 mM in the culture media, unless stated otherwise. When real hypoxia was used, as in Figure 4g, a 3% O2, 5% CO2, balance N2 environment was created by placing the culture dishes in a Modular Incubator Chamber (Embrient, Inc., formerly Billups-Rothenberg, Inc., MIC-101) and flushing the headspace with the desired gas mixture for 3 minutes. A Dual Flow Meter (Embrient, Inc., formerly Billups-Rothenberg, Inc., DFM-3002) was used to achieve the appropriate influx from two different compressed gas cylinders—one cylinder contained 5% CO2 / balance O2 (Airgas, #Z02OX9512000033), and the other contained 5% CO2 / balance N2 (Airgas, #X02NI95C2003071). Cultures were re-transfected with the appropriate recording cassettes every 4 days and re-induced again approximately 16 hours after transfecting.

### Illumina library preparation for the high-throughput NNK library screens

For each of the 3 site saturation mutagenesis libraries, all cells from the end of the *in vivo* assay were subjected to genomic DNA (gDNA) extraction with a QIAamp DNA Mini kit (Qiagen, 51304), then combined into one pool per library. A total mass of 150 µg of the resulting gDNA from libraries 1, 2, and 3, was used as a template for “1^st^ round PCRs,” which amplified the region of interest while simultaneously adding partial Illumina adaptors (Supplementary Table 7). The large mass of gDNA template for the 1^st^ round PCRs was divided across many replicate reactions per library (38-75 reactions per library, depending on how much gDNA could be tolerated in individual PCRs for each library), with 3 different sets of unique multiplexing barcodes on the forward and reverse primers for each library (this split each library into 3 discernable technical replicates). The maximum possible mass of gDNA was used in each PCR, as this allowed us to sample gDNA from many separate cells and reduce the risk of PCR jackpotting. Similarly, each library was amplified in many replicate PCRs to increase the total amount of gDNA that could be sampled, and to dilute the potential impact of PCR jackpotting in any single reaction. Phusion Hot Start Flex polymerase (New England Biolabs, M0535) was used for the 1^st^ round PCR, for 20 cycles of amplification with 62°C, 63°C, and 66°C annealing temperatures for libraries 1, 2, and 3, respectively, and an extension time of 40 seconds. Products from the 1^st^ round PCRs were pooled together for each library, then approximately 1/10^th^ of the volume of each pool was purified with a 0.9:1 ratio of AMPure XP beads:PCR mix (Beckman Coulter, A63881). The complete volume of the AMPure-cleaned 1^st^ round PCR products for each library was used as the templates for a “2^nd^ round PCR,” which added the remainder of the TruSeq HT i5 and i7 Illumina adaptor sequences. The 2^nd^ round PCRs were performed in 10 replicate reactions for each library. Phusion Hot Start Flex amplified the 2^nd^ round PCRs in 20 rounds of thermal cycling with a 63°C annealing temperature and 40 second extension time. The final PCR products for each library were pooled, purified again with 0.9:1 AMPure XP beads:sample, and sequenced with an Illumina NovaSeq6000 using the paired-end 250 protocol at the UC Irvine Genomics, Research, & Technology Hub.

### Illumina library preparation for individual TdT assays and CHYRON experiments

Genomic DNA (gDNA) was extracted from HEK293Ts using the QIAamp DNA Mini kit (Qiagen, 51304). For each sample, approximately 200 ng of the resulting gDNA was used as a template for a single PCR that added sample-specific multiplexing barcodes and partial Illumina adaptors while amplifying the locus of interest (the HEK3 locus or CHYRON locus) (Supplementary Table 7). For this, Phusion Hot Start Flex polymerase (New England Biolabs, M0535) was used according to the manufacturer’s protocol, with an annealing temperature of 58°C, 1 minute extension time, and 30 rounds of thermal cycling. PCR products with unique multiplexing barcodes were pooled together and subsequently purified with AMPure XP beads (Beckman Coulter, A63881), with a 0.9:1 ratio of beads:sample. Purified pools were sequenced by Azenta Life Sciences, with the Amplicon-EZ service. This involved further amplification to constitute the TruSeq HT i5 and i7 adaptors, then sequencing on an Illumina HiSeq 2500 with a paired-end 250 protocol.

### *In vitro* expression and purification of TdTs

T7 express *E. coli* (NEB) was transformed with the final pET28a constructs. Transformation outgrowth was plated on kanamycin and incubated overnight at 30°C. Colonies were selected and grown overnight in 5 mL LB media with 25 µg/mL kanamycin. The next day, these overnight cultures were diluted 400x in 60 mL of fresh LB media and 25 µg/mL kanamycin. The cultures were grown to an OD of 0.4-0.8 (∼3 hours) at 37°C, induced with 1 mM of IPTG, and incubated for 16 hours at 15°C. After 16 hours, the cultures were centrifuged at 4000xg for 20 minutes, and the supernatant was discarded.

To purify TdT, TALON Metal Affinity Resin (Takara) protocols were used, and all steps were performed at 4°C on ice. To lyse the harvested cells, pellets were resuspended in xTractor buffer (20 mL per 1 g pellet) with 1X lysozyme (200 uL of 100X lysozyme solution per 1 g pellet) and DNase I solution (40 uL of 5 units/uL per 1 g pellet). After incubating on a shaking platform for 20 minutes at 175 RPM, the tubes were centrifuged at 12,000xg for 20 minutes. The supernatant (soluble fraction) was applied to 100 uL of equilibrated TALON Resin and incubated on a shaking platform for 20 minutes to allow the His-tagged TdT to bind the resin. Thereafter, the resin was pelleted by centrifugation (700xg for 5 minutes at 4°C) and the supernatant was carefully removed. The resin was washed twice by incubating it with 1 mL equilibration buffer for 10 minutes (Takara), centrifuging the resin (700xg for 5 minutes) and discarding the supernatant. After washing, the resin was resuspended in 100 uL of equilibration buffer, briefly vortexed, and transferred to a 2 mL gravity column (Takara). The end-cap of the column was removed to drain the buffer and 500 uL of wash buffer was added to wash the resin once more. 500 uL of Elution Buffer (Takara) was then added to the column to elute the TdT into a clean 1.5 mL tube. The protein concentration was measured using a Nanodrop (absorbance at 280nm). TdT expression and purity was verified and analyzed via SDS-PAGE. The elution was buffer-exchanged and concentrated using 10K Amicon-Ultra 0.5 centrifugal filters (10,000 MWCO) into TdT storage buffer (200 mM KH2PO4 and 100 mM NaCl at pH 6.5). The fractions were aliquoted and stored at –80°C.

### *In vitro* TdT extension assays

All TdT extension reactions were set up on ice and contained final concentrations of 0.1 uM of common sequence 1 primer (CS1_5N, 5’-ACACTGACGACATGGTTCTACANNNNN-3’), which has 5 degenerate bases at the 3’ end of the primer. 400 uM of dNTP mix (100 uM of each), 1X NEB TdT reaction buffer, purified TdT enzyme (varying amounts depending on the variant), and nuclease-free water were added to 50 uL. The reactions were performed for 1 hour at 37°C.

For each variant, a range of TdT concentrations were tested to titrate TdT such that the approximate average extension lengths after 1 hour at 37°C were in the appropriate length range for next-generation sequencing. We aimed for an average extension length below ∼60nt. For these titration experiments, the extension products were analyzed by urea-PAGE. A FAM-labelled version of the CS1_5N primer was used to enable this. 8uL of the completed extension reaction was mixed with 12 uL of BioRad 2x TBE urea sample buffer and boiled for 10 min. The samples were loaded into a 10% polyacrylamide TBE urea gel (bioRad 4566036) and run at 200 V for 40 min. The gels were imaged on a Sapphire Biomolecular Imager using 100 um pixel size and λex = 488 nm and λem = 520 nm. The imaging gain was adjusted for each experiment to avoid saturation. The final concentrations found to be appropriate for extension reactions to be sequenced *via* NGS are shown in the table below.

| TdT variant | TdT yield following purification (mg/mL) | Final TdT stock concentration following buffer exchange (mg/mL) | Final TdT concentration in extension reaction for NGS (µg TdT/50uL reaction) |
| --- | --- | --- | --- |
| Wild-type TdT | 0.26 | 0.44 | 0.2 |
| Unbiased1 (MQFCRDLR) | 0.14 | 0.31 | 0.2 |
| Unbiased2 (RAFD) | 0.18 | 0.41 | 0.2 |
| Unbiased3 (KQFRRDLR) | 0.31 | 0.47 | 0.3 |
| A/T-biased1 (LQFKRDLL) | 0.21 | 0.25 | 0.1 |
| A/T-biased2 (IQFKRDLA) | 0.27 | 0.38 | 0.1 |
| G/C-biased1 (DQFDRDLR + DAMD) | 0.27 | 0.31 | 0.3 |
| Wild-type-like (DLY) | 0.25 | 0.40 | 0.2 |

### Next-Generation Sequencing of *in vitro*-extended samples

To prepare the extension reactions for NGS, common sequence 2 (CS2, 5’Phos-AGACCAAGTCTCTGCTACCGTA-3’) primer was first ligated to the 3’ end of the TdT-synthesized polynucleotides. 2 uL of the extension reaction product was used in a ligation reaction containing final concentrations of 1 uM CS2 primer, 10 units of T4 RNA ligase (NEB), 1X T4 RNA ligase reaction buffer (NEB), 10 uL of PEG 8000 (NEB), 1 mM of ATP, and nuclease-free water up to 20 uL reaction volume. The ligation reaction was carried out at 25°C for 2 hours.

PCR was performed to prepare the ligation products as double-stranded, barcoded samples ready for sequencing according to previous protocols^71^. Primer sets from the Access Array Barcode Library for Illumina Sequencers (Fluidigm) were used for barcoding. The Fluidigm primer sets contain the complement of CS1 (the defined sequence of CS1_5N) as the binding target of the forward PCR primer and the complement of CS2 as the reverse primer. The Fluidigm primers also contain a unique barcode in the reverse primer and Illumina adapters in the forward and reverse primers for sequencing. The PCRs consisted of 2 uL of the ligation product, 1X Phusion High-Fidelity PCR Master Mix with HF Buffer (NEB), and 400 nM forward and reverse Fluidigm PCR primers in a 20 uL reaction volume. Products were initially denatured for 30 seconds at 98°C, followed by 20 cycles of 10 seconds at 98°C (denaturation), 30 seconds at 60°C (annealing), and 30 seconds at 72°C (extension). Final extensions were performed at 72°C for 10 minutes. PCR samples were sent for sequencing at Rush University’s Genomics and Microbiome Core Facility. Quality control (QC) and Illumina MiniSeq protocols were the same as described in previous studies^71^.

### Computational analysis of NGS data

All analyses were conducted with custom Python scripts, which will be made available at github.com/liusynevolab/TdT_high-throughput_screen.

For the high-throughput screen, the amplicons that were sequenced were ∼1 kb long (sufficiently long to cover the NNK codons in the TdT gene near the 5’ end of the amplicons, and the Cas9 target site near the 3’ end), so the Illumina reads (paired end 250) did not cover the full length of the amplicons. Thus, forward and reverse reads for each molecule were simply concatenated, such that the NNK sequences from a given read were stored in the same string as the TdT-mediated insertion at the Cas9 target site. In other words, nothing of importance was in the middle of the ∼1 kb amplicons, and there was no sequence overlap between the forward and reverse reads, so there was no need to align them.

After concatenation, reads of incorrect amplicons (*e.g.*, primer dimers or non-specific amplicons) were removed. Extremely short amplicons, such as primer dimers, were identifiable without performing a formal sequence alignment because they were marked with long repetitive stretches of “G” in their reads. This is because Illumina uses sequencing by synthesis with a fluorescent readout to indicate which base is incorporated in each round of synthesis, short molecules prematurely stop generating fluorescence in the late rounds of sequencing by synthesis, and lack of fluorescence is miscalled as guanine by the Novaseq basecalling software. Thus, any reads with homopolymeric stretches longer than 15 bp were assumed to be primer dimer and removed. 15 bp is the maximum insertion size we attribute to one round of TdT insertion mutagenesis (anything larger tends to be due to homologous recombination at the Cas9 cut site), so this cutoff would tolerate even the highly unlikely hypothetical case of an extremely biased TdT inserting up to 15 bp of a single nucleotide.

Once the non-specific short reads were filtered out, the remaining reads were demultiplexed according to the barcodes on the Illumina sequencing primers. Each library was labeled with 3 different pairs of multiplexing barcodes, so a total of 9 barcode pairs were expected across the 3 libraries. Any reads that had unexpected combinations of barcodes on the 5’ and 3’ ends of the read were assumed to be the product of template-switching during library preparation and were therefore discarded.

The demultiplexed reads were aligned to an appropriate reference sequence for each library, using Mapp as in Perli et al^58^. Alignments are represented by Mapp as strings containing “M” for match, “X” for mismatch, “D” for deletion, and “I” for insertion at each base in the read. These alignments were used for a quality filtering step, wherein reads that had >50 mismatches (10% of bases in the read were miscalled), >250 deletions (half of the read is deleted), or >15 contiguous insertions (too large to be a TdT-mediated insertion) were removed.

The DXMI characters were used to extract NNK sequences and insertion sequences from the aligned reads. Codons that were mutagenized in each library were designated with “NNN” in the reference sequence, so the alignment would always indicate three mismatches (“XXX”) when aligning reads with the randomized codons. The analysis script searched for the expected alignment at the sites of saturation mutagenesis and checked for 2 perfectly aligned bases (“MM”) on either side of the mutated residues (“MMXXXMM”). Similarly, the DXMI characters were used for extracting TdT-mediated insertion sequences and NHEJ-generated deletions from the Cas9 cut site in each read. A small window around the expected location of the Cas9 cut site was checked for the presence of an “I” or “D” in the Mapp alignment. If an “I” was present, the adjacent alignment characters were scanned for additional, contiguous “I’s”, so the full length of each insertion was captured. Once the entire string of “I’s” was found, the analysis script ascertained that the read has perfectly aligned bases on either side of the insertion (“MM I_n_ MM”), and if so, extracted the insertion sequence. Any Cas9 target site that harbored one or more “D(s)” at the Cas9 target site was counted as a deletion, even if accompanied by “I(s)”. The sequences at each Cas9 cut site were binned according to which TdT mutant gene (NNK codons) were linked on the same read. Ultimately, each TdT variant has a collection of hundreds to thousands of editing outcomes associated with it.

Having binned all insertions and tallied all deletions for each TdT variant, the script quantified the following metrics for each variant: total number of extracted insertion sequences, average length of insertions, average A/T bias among all inserted nucleotides, average A/T bias for insertions of each observed length, number of insertion sequences of each length, “insertion fraction” (*i.e.*, fraction of reads with an insertion / reads with any indel), and number of unique insertion sequences for each TdT variant. For each library, a comprehensive Pandas dataframe was created, reporting all of these metrics for each TdT variant. Microsoft Excel was used to manually inspect the dataframe, sorted by overall A/T bias of each TdT variant, and select variants of interest for follow-up experiments. These comprehensive dataframes for each library are made available in Supplementary Tables 1-3. Matplotlib was used to create histograms of insertion fractions and scatterplots of A/T bias vs. average insertion length for each TdT variant.

For individual assays of TdT variants and CHYRON experiments, samples were demultiplexed *via* sample-specific barcodes on the forward and reverse reads. Insertions were extracted by checking each read for the expected 15-bp sequence upstream of the Cas9 target site (within a user-defined Hamming distance), then scanning for the 15-bp sequence downstream of the Cas9 target site (within a user-defined Hamming distance), and finally grabbing the intervening sequence. If this initial alignment failed to locate the Cas9 target site (*e.g.*, because the locus was partially deleted, a normal editing outcome for Cas9 cutting), it searched for an alignment with 15-bp sequences further out from the cut site. The distance between the upstream and downstream aligned sequences was used to calculate the size of deletions. The lengths of all insertions and deletions were tallied, to create a histogram of indel sizes. Compositions of the extracted insertions (average insertion length, overall A/T content across every inserted nucleotide, and average incidence of all 4 natural nucleotides for each observed insertion length) were reported.

### Analysis of amino acid compositions among differently-biased TdT library members

Our goal here was to assess the enrichment of particular amino acids at the randomized positions among library members with altered nucleotide insertion biases. Altered-bias TdTs were included in this analysis only if they produced at least 250 reads containing insertions (to ensure the calculated biases were reliable) and had insertion fractions of 0.7 or higher (because inactive variants are not practically useful). Therefore, to define the baseline composition of amino acids from which enrichment could be calculated, we assessed the compositions of residues among all library members that were observed in enough instances in the NGS data to ultimately have had a chance of passing these filters. In other words, we assessed the composition of TdT variants that were present in at least N = 447 instances in the NGS data because anything less would result in the generation of fewer than 250 insertions at a minimum insertion fraction of 0.7 and a typical Cas9 DSB break generation efficiency of 80% (*i.e.,* if N < 447, then 447 * 0.7 * 80% < 250). The TdT variants that were present in at least 447 instances in the NGS data were used to generate the baseline composition of amino acids for the TdT libraries (Supplementary Figure 1). From the baseline composition of amino acids, the enrichment of residues was calculated for the TdTs that were sorted into bias bins (Figure 1d) as:

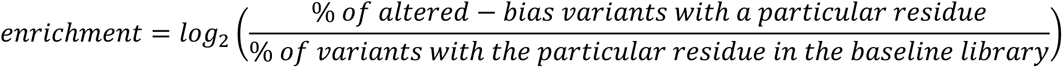

### Considerations informing experimental parameters of multiplex assay execution and analysis

Each TdT library was constructed using NNK-codon randomization at 3 positions (N = any nucleotide, K = G or T). Therefore, the theoretical diversity of DNA sequences in each library was (4*4*2)^3^ = 32,768 sequences. In choosing experimental parameters during library cloning steps, we aimed for 12-fold coverage because this created a high probability (99% likelihood) that 100% of the theoretical diversity would be represented^68^. In the end, each library’s theoretical diversity was covered 70.7-fold to 80-fold (Supplementary Table 8) such that the probability of excluding any library member was vanishingly small. The hp-minipreps encoding the libraries were assumed to be more than sufficient to cover the theoretical diversity of libraries, since a single nanogram of our 10.3 kb library plasmid contains approximately 9.5E+07 molecules (3000-fold library coverage). This abundantly diverse pool of DNA was used for lentivirus production, achieving titers of 1.6E+08, 8.5E+08, and 8.4E+08 infectious particles / mL, for libraries 1, 2, and 3, respectively. At least 100 uL of each virus was provided by the UCSF ViraCore, constituting approximately 500-fold coverage. During transduction, we aimed for a minimum of 12-fold coverage of each library’s theoretical diversity, and we exceeded this goal as much as feasible. Ultimately, 3x T-150 flasks at ∼30% confluency were transduced at a MOI of 0.3 for each library. Assuming 3,300 cells / mm^2^ in a confluent culture dish (empirically approximated using a hemacytometer), each transduction produced nearly 1E+07 infected cells (304-fold library coverage). In short, throughout all steps leading to establishment of TdT libraries in HEK293T cells, high coverage of each library’s theoretical diversity was maintained.

After selecting for transduced cells, our goal was to not only represent every library member in the HEK293T cell population, but to have every library member represented in enough cells in the population so as to gain confident estimates of the behavior of each TdT variant. Since each cell containing a given TdT variant represented an independent trial of the variant’s insertional behavior when the assay is initiated through the transfection of Cas9 to generate the DSB, we were interested in estimating how many cells per TdT variant we would need to gain confident estimates on insertion bias. We considered the simple case where each nt that TdT adds is an independent event with binary outcomes: an A/T insertion or a G/C insertion. The probability, Pr, of having exactly *k* nts that are A/T in *n* trials is:

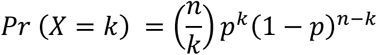

for a library member with an A/T bias of *p*. This probability function was used to calculate the fiducial limits of *p* given an observed *k* / *n*, with the Clopper-Pearson method^72^. We assessed the 95% confidence intervals for different levels of A/T bias (*p*) as a function of the number of trials (*n*). We settled on a target of 250 trials per library member to strike the right balance between being practically feasible and providing an acceptable degree of certainty in distinguishing large differences in bias measured for different variants. For example, at 250 trials, the 95% confidence intervals for the true A/T biases, given an observed % A/T, are shown below:

| Observed % A/T in TdT-mediated insertions (i.e., $k$ successes / $n$ trials) | Actual value of $p$ (i.e., A/T bias), 95% confidence intervals |
| --- | --- |
| 10% | 5-15% |
| 50% | 44-56% |
| 90% | 85-95% |

Most TdT variants were expected to make longer than 1 bp insertions on average (wild-type insertions are ∼3 bp). However, we considered the conservative case that an active TdT variant would only insert one nt at the Cas9-generated DSB. Therefore, we aimed to observe at least 250 insertions per TdT variant in our NGS dataset when making experimental choices for the multiplex assay. In particular, the goal of achieving 250 insertions per TdT variant determined how many cells we should transfect with Cas9 and its sgRNA to initiate TdT insertions. Given that each library had 32,768 variants, that we wanted to have ∼250 independent trials for each, that the efficiency of Cas9 and its sgRNA in DSB generation is ∼80% when transfected, and that the frequency with which TdT usually makes an insertion at a Cas9-generated DSB is approximately ∼75%, we wanted to transfect ∼1.4E+07 cells (*i.e.*, (32,768 * 250) / (0.8 * 0.75)).

3x T-150 flasks, each approximately 25% confluent, were transfected for each library. This amounts to approximately 1.25E+07 cells per flask at the time of transfection. We assumed Cas9 would be expressed and start cleaving its target locus at 24 hours post-transfection, at which point the cultures would have 2.5E+07 cells, ∼80% of which would generate a DSB through the action of Cas9 and 75% of which would receive a TdT insertion. The cultures were incubated for a total of 72 hours after transfection, which is long enough to allow Cas9 cutting, TdT-mediated insertion, and DNA repair, but short enough to prevent excessive outgrowth. Each cell should have divided only 1-2 times after acquiring a TdT-mediated insertion. All cells in a given library were pooled together upon collection, and the genomic DNA (gDNA) from the complete pool of HEK293Ts for each library was extracted.

To faithfully capture the insertion outcomes in the gDNA, we took precautions to PCR from as much of the gDNA as possible to avoid amplification biases. Ultimately, we used 150 µg genomic DNA as template for each library (divided across many smaller PCRs). Assuming 8 pg gDNA per HEK293T cell, this corresponds to gDNA from 1.9E+07 cells in the PCRs for Illumina library prep. The number of PCR amplification cycles was also minimized during library prep – we found 20 cycles to be sufficient for both the “1^st^ round” and “2^nd^ round” PCRs. Final Illumina libraries were sequenced at 12.6-fold coverage of the desired 250 insertions per variant, for a total of 3.1E+08 reads. Because our Illumina libraries had a larger molecular weight (∼1 kb) than typical Illumina libraries (<500 bp), our target amplicons clustered somewhat inefficiently on the NovaSeq flow cell. After filtering out primer dimers and low-quality reads, we retained 1.0E+08 Illumina reads, and we extracted averages of 175, 215, and 367 insertion sequence reads per TdT variant (variants defined by DNA sequences) for libraries 1, 2, and 3, respectively. When synonymous TdT variants were binned together, we had averages of 1361, 1680, and 2816 insertion sequence reads per TdT variant (variants defined by amino acid sequences).

Although we observed many insertion sequence reads per TdT variant for each of the libraries, these reads are not all the result of independent TdT insertion events. As stated above, 2.5E+07 cells * 80% * 75% = 1.5E+07 cells were estimated to have generated a DSB by Cas9 and received an independent insertion through TdT activity. After TdT insertions occurred, these cells divided such that each independent TdT insertion event was represented a few times in the resulting population of cells. Then, gDNA from 1.9E+07 cells was used for NGS library prep with roughly 80% * 75% of these gDNA molecules (*i.e*., 1.14E+07) containing insertions based on our above estimates of Cas9 DSB generation and TdT insertion fraction. From these numbers, we can roughly estimate how many independent TdT insertion events are represented in the estimated 1.14E+07 gDNAs with insertions. The probability, *P*, that any particular outcome in the 1.5E+07 independent TdT insertion outcomes is not sampled in 1.14E+07 gDNAs is:

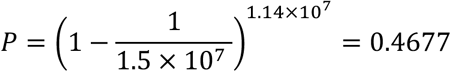

The probability that any particular outcome in the 1.5E+07 independent TdT insertion outcomes is sampled at least once in the estimated 1.14E+07 insertion-containing gDNAs is therefore:

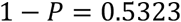

This means that 53.23% of the 1.14E+07 insertion-containing gDNAs per library represents independent TdT insertions.

The number of TdT variants in each library is no more than 32,768 such that each library variant is represented, among the 1.14E+07 insertion-containing samples, 348 times on average. Roughly 53% of those 348 instances of each TdT variant’s insertion outcomes should be independent, meaning that an average of no more than ∼184 reads per TdT variant (variants defined by DNA sequence) for libraries 1, 2, and 3, respectively, correspond to instances of independent TdT insertions. ∼184 independent insertion observations should allow for confident bias determination, especially given that each insertion contains multiple nts on average.

## Acknowledgements

We thank all members of the Liu lab for helpful discussions throughout this work. We thank Gordon Rix for technical assistance with python programming, and for making Pymol images. Melanie Oakes provided insightful discussion that was pivotal in planning Illumina sequencing for the high-throughput screen. This work was made possible, in part, by the Genomics, Research, and Technology Hub at the University of California, Irvine and the High Performance Community Computing Cluster at the University of California, Irvine. This work was funded by NIH 1R35GM136297 to CCL and AHA Predoctoral and NSF Graduate Research fellowships to CKC. We thank Hana El-Samad, Keith Joung, Randall Moon, Yvonne Chen, Eric Campeau, and Paul Kaufman for plasmids. Next-generation sequencing data for *in vitro* TdT activity measurements were generated with the help of the Rush Genomics and Microbiome Core Facility.

## Author Contributions

CKC, TBL, PIK, JM, and CCL selected residues to target for saturation mutagenesis. CKC, TBL, MM, and CCL designed experiments. CKC performed all experimental work, except for the *in vitro* assays and corresponding NGS preparation, which were performed by MM. CKC, TBL, MM, and CCL analyzed data. CKC and MM wrote code for analyses. CKC and CCL wrote the manuscript with input from all other authors. KEJT and CCL procured funding and oversaw the project.

## Supplementary Information

**Supplementary Figure 1.**
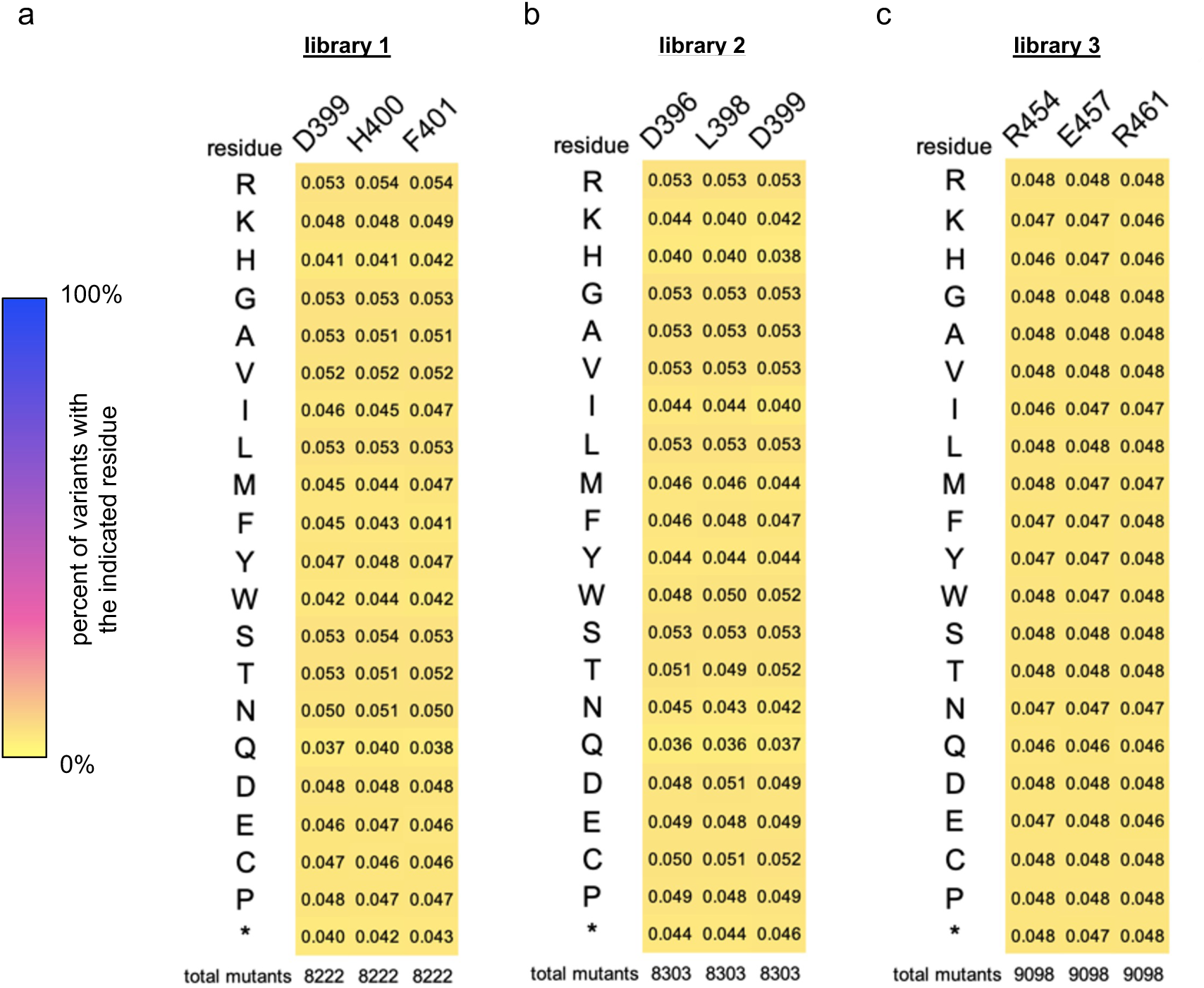
Heatmaps displaying the compositions of amino acids at the target sites for saturation mutagenesis (NNK loci) in the full TdT libraries. **a-c,** The amino acid compositions were plotted for libraries 1-3, for all TdT variants that were sequenced in enough instances to have a chance of passing the activity and insertion count cutoffs (see Methods). Enrichment of amino acids in specialized bias bins (in Figure 1d) were calculated relative to these values.

**Supplementary Figure 2.**
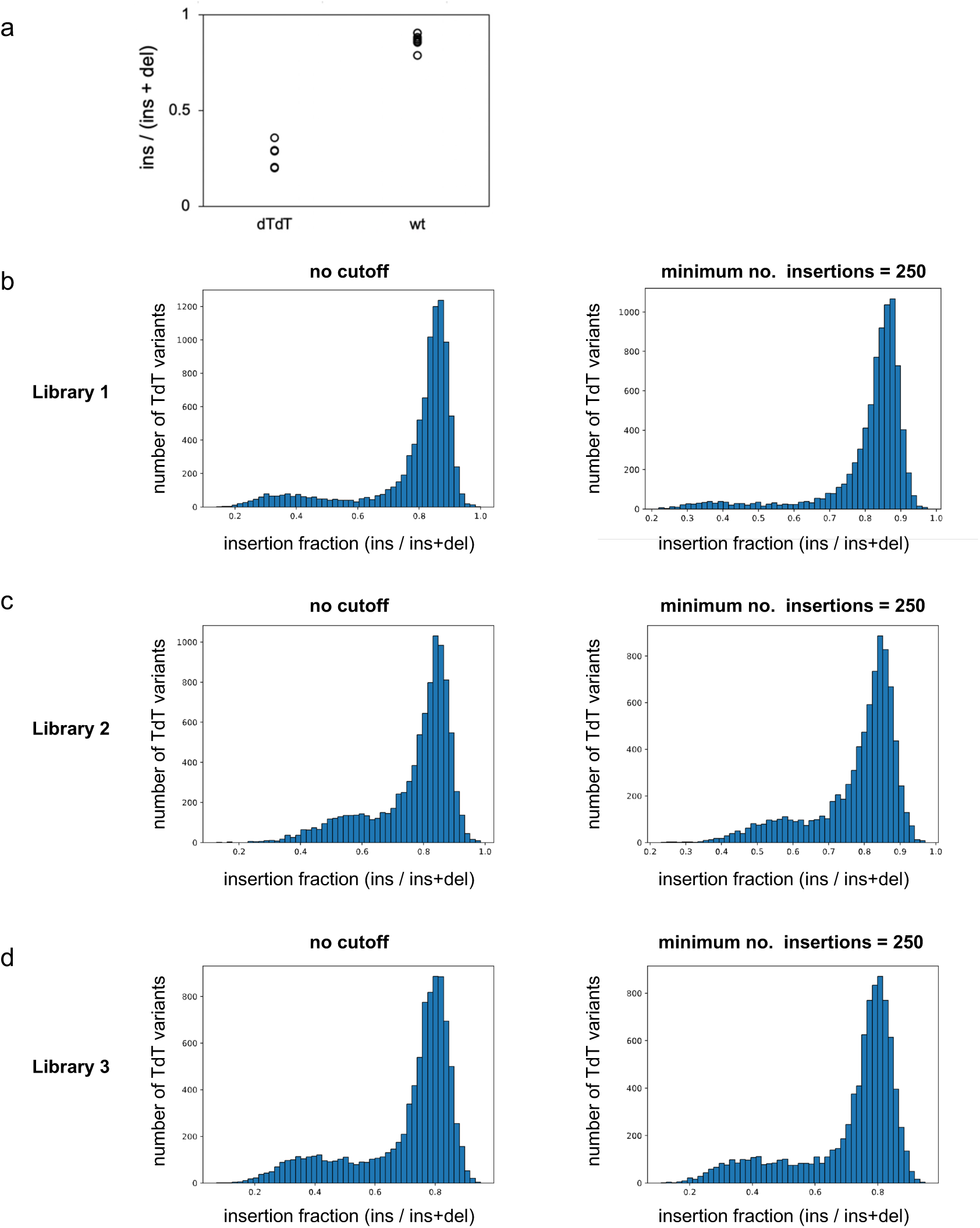
Assessment of library members’ level of activity. **a**, The frequency of insertions relative to any indel (”insertion fraction”) is shown for active TdT (wild-type, wt) and a catalytically dead variant (dTdT). Five and eight biological replicates are shown, respectively, for dTdT and wt TdT transiently transfected into HEK293T cells with Cas9 targeting the HEK3 genomic locus. **b-d,** For libraries 1 (base of Loop1 NNKs), 2 (mid Loop1 NNKs), and 3 (alpha helix NNKs), histograms are shown for the insertion fractions of the library members. The raw data for every TdT variant (no cutoff), and the data after filtering out mutants that were sampled with low statistical power (minimum number of insertions = 250) are shown. Each TdT mutant corresponds to translated protein sequences, such that synonymous mutants in the library are counted as a single mutant.

**Supplementary Figure 3.**
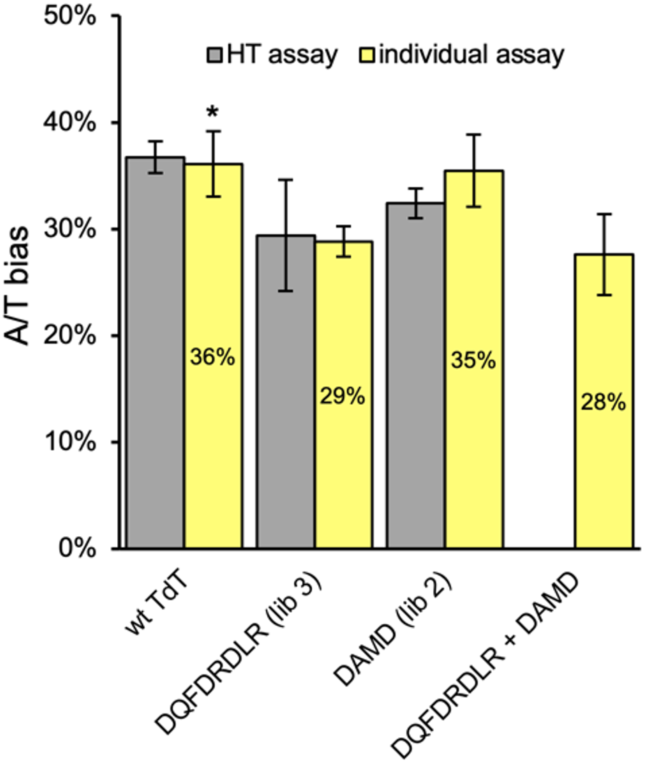
Observed nucleotide biases of two G/C-biased TdT variants that were selected from the high-throughput (HT) assay, and a child variant that combines mutations from both. The amino acid sequences spanning all randomized residues for each variant are indicated (DQFDRDLR from library 3, residues 454-461; DAMD from library 2, residues 396-399). The double mutant (DQFDRDLR + DAMD) was used for subsequent *in vitro* characterization and *in vivo* DNA recording experiments. For the HT assay data, the average of 3 technical replicates is plotted, with error bars showing the standard deviation. For individual assays, unless stated otherwise, the average of 3 biological replicates is plotted, with error bars demarcating the standard deviation. * = 7 biological replicates are shown for wildtype TdT assayed individually.

**Supplementary Figure 4.**
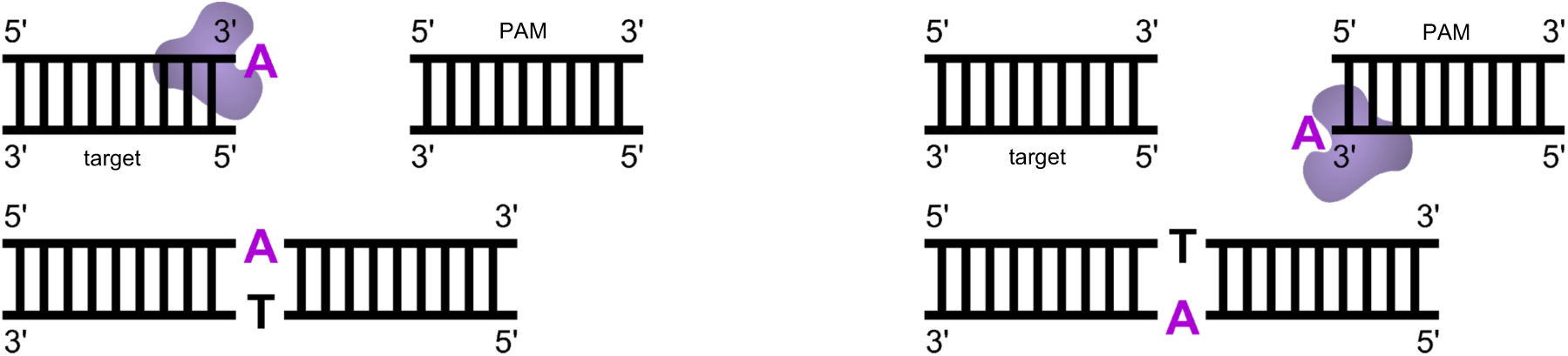
Illustration of ambiguous insertion identities for complementary bases at double-strand breaks (DSBs). After Cas9 induces a DSB, TdT has two available primer strands to extend. If TdT uses the top strand (PAM-containing strand) as its primer, the inserted nucleotide will be read out directly during NGS. Alternatively, if TdT adds a nucleotide to the bottom strand (Cas9’s target strand), the complementary base will be added to the top strand during DNA repair, and subsequently read out during NGS. Thus, TdT’s preference for A vs. T and C vs. G cannot be determined from *in vivo* assays at DSBs, but bias for A/T vs. G/C can still be accurately measured.

